# Sequence determinants of the hypomobility of intrinsically disordered proteins in SDS-PAGE

**DOI:** 10.64898/2026.03.24.714011

**Authors:** Ankush Garg, Maciej B. Gielnik, Magnus Kjaergaard

**Author notes:** equal contribution (Ankush Garg, Maciej B. Gielnik).

## Abstract

Proteins with intrinsically disordered regions (IDRs) migrate at a higher apparent molecular weight in sodium dodecyl sulfate-polyacrylamide gel electrophoresis (SDS-PAGE) complicating their analysis and identification. Here, we investigate the sequence determinants of the hypomobility of IDRs using a series of synthetic low complexity domains. We find that negative charge increases the apparent molecular weight, but neutral polar tracts also have abnormally slow migration. Positive charge and hydrophobic residues decrease the apparent molecular weight, although lysine residues show a biphasic effect with decreased migration at high fractional contents. Combinations of residues show that different sequence contributions to the apparent molecular weight are not additive. The results can be rationalized by the protein-decorated micelle model by considering both SDS binding and the compaction of protein SDS-complexes.

## Introduction

About a third of eukaryotic proteins contain at least one intrinsically disordered region (IDR) – a region without a well-defined three-dimensional structure.^1^ Soon after intrinsically disordered proteins (IDPs) were recognized as a general phenomenon, it was noticed that they behave differently than folded proteins in SDS-PAGE.^2^ The SDS-PAGE assessed molecular weight of an IDP can be twice its actual molecular weight, although this ratio varies between proteins, which can cause practical problems during protein purification.

IDRs are enriched in polar and charge residues at the expense of hydrophobic residues,^3,4^ which is believed to cause their abnormal SDS-PAGE behaviour. The reverse occurs in membrane proteins that often migrate at lower apparent molecular weights (M_w,app_) due to their hydrophobicity.^5^ In soluble proteins, SDS-PAGE hypomobility has been linked to high negative charge. Point mutations in SOD1 shift SDS-PAGE mobility primarily when they target a negatively charge cluster.^6^ In a set a truncation variants, the fraction of negatively charged residues correlated strongly with hypomobility - in contrast to other sequence features - which led to a proposed linear relationship between the M_w,app_ per residue and the fraction of negative residues.^7^

In SDS-PAGE, electrophoretic mobility is determined by the net negative charge - primarily from bound SDS – that drives proteins towards the anode, and the friction in the gel determined by the size and shape of the protein. The dimensions of chemically denatured proteins and IDPs both follow power law dependences on chain length_^8,9^_ suggesting that chain compaction alone is not sufficient to explain the different mobility. Protein-SDS complexes are best described by the *protein-decorated micelle* model, where extended protein chains wrap around and insert into SDS micelles.^10^ NMR studies of the intrinsically disordered α-synuclein shows that SDS micelles mainly interacts with the hydrophobic segments leaving the negatively charge C-terminus unperturbed.^11^ Molecular dynamics simulations suggest hydrophobic residues from α-synuclein insert into the SDS micelle, positively charged residues form electrostatic interactions the SDS head groups, and negatively charged residues protrude into the surrounding solvent.^12^ Protein-SDS complexes thus have sequence-specific effects that might translate into different mobility during SDS-PAGE.

It is still largely unpredictable at which apparent molecular weight an IDR will migrate in SDS-PAGE, likely due to the absence of systematic studies. Here, we investigated how sequence features of IDRs affect apparent molecular weight assessed by SDS-PAGE. We used a host-guest approach, where different residues are titrated into a featureless IDR. We found that negatively charged amino increase M_w,app_ strongly, but neutral polar tracts also have an elevated M_w,app_. Positively charged residues have a bi-phasic effect, initially decreasing and then decreasing M_w,app_. Hydrophobicity reduces the apparent molecular weight in uncharged proteins but do not counteract the effect of charge. In combination, our results support a model where high M_w,app_ of IDRs derive from the dynamic structures formed in complex with SDS micelles.

## Results & Discussion

Inspired by an IDR biosensor,^13^ we expressed synthetic IDRs as a loop between two interacting coiled-coils with orthogonal terminal purification tags (Fig. 1a). The IDR was flanked by thrombin cleavage sites to allow release of the IDR for other studies. Many of the isolated IDRs were faint or not visible on stained gels – even when size exclusion chromatography showed that the free IDR was present (Fig. S1). The failure to detect these IDRs is likely due to a lack of fixation, a lack of dye staining, or both. Anecdotal evidence suggests that this is common for small IDRs. Therefore, we analysed M_w,app_ in the expression construct by subtraction M_w,app_ of a construct without an IDR (Fig. 1a,b). This result in the M_w,app_ of the IDR as reported by SDS-PAGE, ΔM_w,app_.

**Figure 1:**
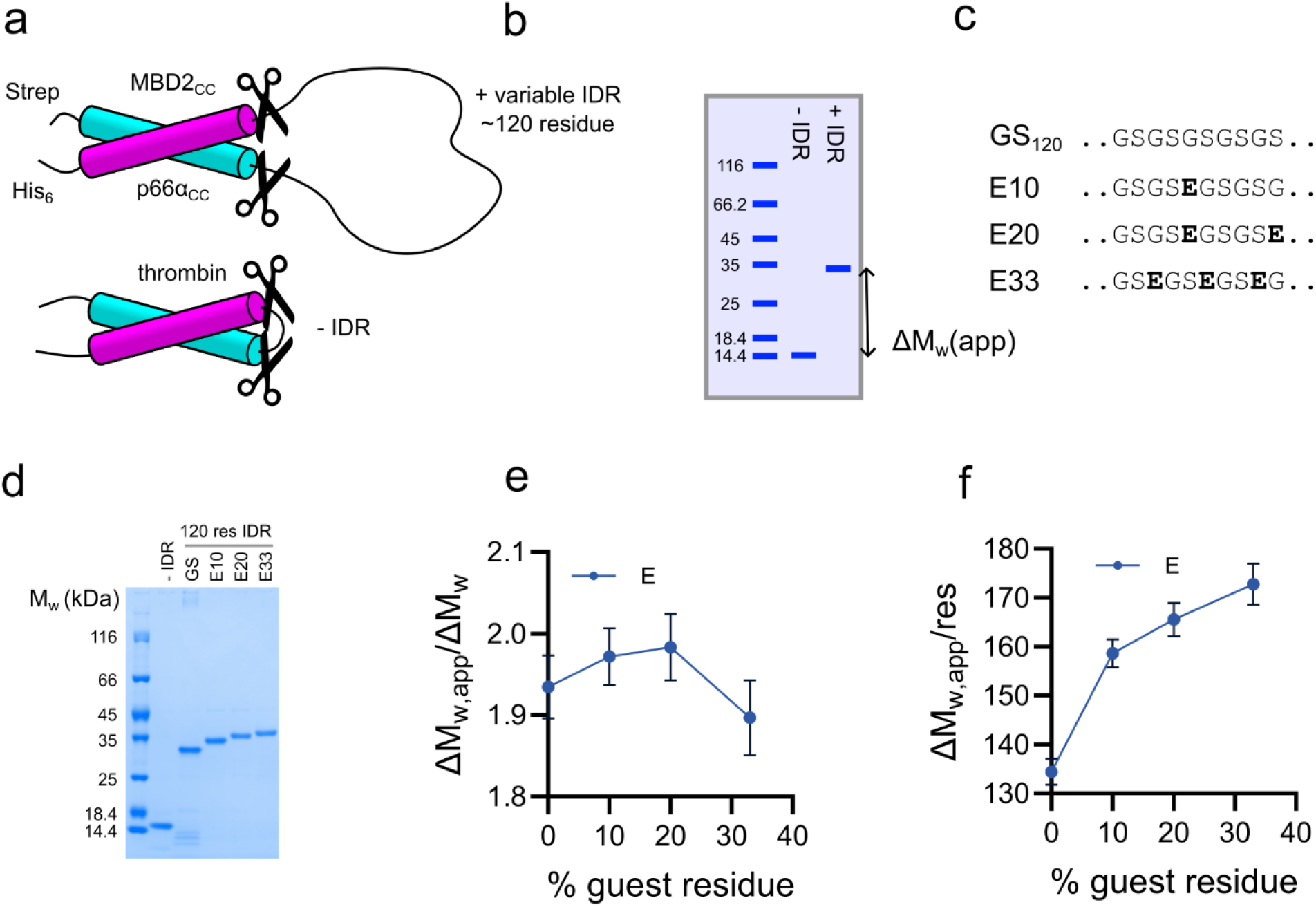
Assessing M_w,app_ of an IDR expressed as a loop. a) Schematic representation of the fusion proteins used, where IDRs are inserted as loops between two interaction coiled coils. b) The apparent molecular weight of the IDR (ΔM_w,app_) is assessed by SDS-PAGE by subtraction of M_w,app_ of a construct with a minimal loop. c) Sequence variations of the IDR are achieved by titrating in uniformly spaced guest residues in a GS-repeat background. The nomenclature used indicates the guest residue followed by its fractional content, i.e. E10 has 10% glutamate residues. d) Representative SDS-PAGE gel showing the shift in M_w,app_ with increasing content of glutamate residues. e) ΔM_w,app_ of the IDR divided by its theoretical M_w_. F) Apparent molecular weight of the IDR per residue. Error bars represent s.d. (n=3).

We measured ΔM_w,app_ for IDRs of a similar length (~120 residues) but with different chemical properties. We started from a featureless polar tract (GS-repeats) and titrated in uniformly distributed residues to change the overall chemical properties of the IDR (Fig. 1c). In the following, we use a nomenclature where E10 refers to 10% uniformly distributed glutamate residues in GS-repeat background. The GS-repeat IDR has ΔM_w,app_ of 16.2 kDa despite having a theoretical ΔM_w_ of ~8.6 kDa. Titration up to 33% content of glutamate residues increased ΔM_w,app_ to 20.8 kDa (Fig. 1d) suggesting that negative charge contributes to the SDS-PAGE hypomobility as shown previously.^7^ When ΔM_w,app_ was normalized to the theoretical ΔM_w_, the uniform increase observed in the gel reversed at high fractions of glutamate (Fig. 1e). This is a quirk of the GS-baseline, which has a M_w_ of 72 Da per residue compared to the average of 110 Da for most proteins. Any substitution thus increases the theoretical M_w_, which cause the inversion in Fig. 1e. Electrophoretic mobility likely depends on chain length rather than M_w_ *per se*. Therefore, we report the ΔM_w_ per residue (Fig. 1f), which mirrors the visually observed mobility and previously reported trends.^7^

Next, we titrated in residues expected to affect IDP compaction and SDS-binding including charged, aliphatic, and aromatic residues as well as proline (Fig. 2a). High contents of hydrophobic and arginine residues resulted in toxicity or aggregation, but the titrations were continued as far as possible. For all residue types, 10% content was possible although even this low content was challenging for W10 as reflected in a lower purity (Fig. 2b). D10 and E10 have identical ΔM_w,app_ (Fig. 2c) suggesting that the negatively charged residues have similar effects. Positively charged residues lead to an initial decrease in ΔM_w,app_ followed by a gradual increase for lysine. The stronger effect of arginine mirror a myriad of studies showing that the two positively charged residues have distinct effects on IDP compaction and interactions.^14,15^ The bi-phasic effect of addition of lysine residues suggests that it affects M_w,app_ in at least two different ways that dominate at different fractions of residues.

**Figure 2:**
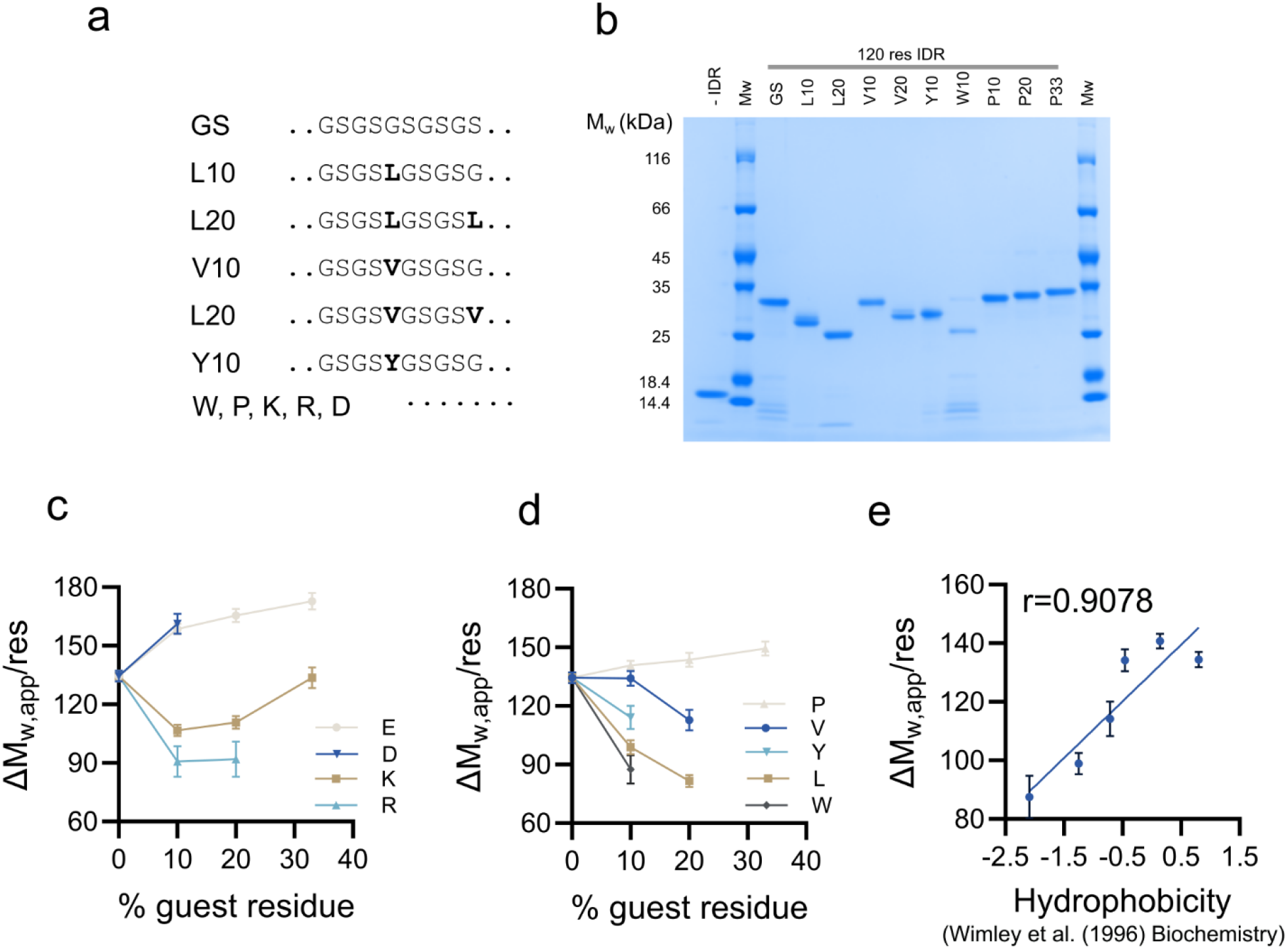
Effect of individual host residues on ΔM_w,app_. a) Sequence patterns used to assess the effect of charged and hydrophobic amino acids on ΔM_w,app_. b) Representative SDS-PAGE used to determine ΔM_w,app_. The M_w,app_ refers to the main band in the gel. c) ΔM_w,app_ per residues for charged residues. d) ΔM_w,app_ per residues for hydrophobic residues. e) Correlation between ΔM_w,app_ per residue of the uncharged guest residue and its hydrophobicity assessed by solvent partition.^17^ The GS-repeat is plotted at the average of G and S. Error bars represent s.d. (n=3).

The four hydrophobic residues all decreased ΔM_w,app_ of the IDR with an apparently linear dependence on fractional content in the limited range permitted by solubility (Fig. 2b,d). In contrast, proline residues increased the ΔM_w,app_ slightly. Proline is not particularly hydrophobic, but tend to expand IDRs.^16^ The apparent M_w_ of IDRs with 10% uncharged residue show a strong correlation (Fig. 2e, Pearson r=0.92) with hydrophobicity assessed from octanol-water partitioning of host-guest peptides.^17^ This is consistent with the finding that the low ΔM_w,app_ of integral membrane proteins is due to their high hydrophobicity and increased SDS binding.^5^

We investigated the additivity of physicochemical features by combining properties with a strong impact on M_w,app_. Simple additivity suggests that combining residues that increase and decrease M_w,app_ would lead these to cancel, whereas combining features that lower M_w,app_ would lead to a larger decrease (Fig. 3a). Alternatively, one physicochemical factor could dominate over the others, leading to an SDS-mobility that follows the mobility of one of the individual factors (Fig. 3a). We tested IDRs with similar fractions of two guest residues and compared their ΔM_w,app_ to IDRs with the same content of individual residue types (Fig. 3b). We combined Glu with Leu, Lys, Val and Tyr, and in each case found that the mobility matched that of a similar content of Glu alone (Fig. 3c-f). This suggested that negative charge dominates over other features that lead to increased M_w,app_ in agreement with previous reports focusing on negatively charged proteins.^7^ We tested the additivity of features reducing M_w,app_ by combining positively charged and hydrophobic residues (Fig. 3g-j). These combinations follow the trend of the positively charged residue, which in most cases resembles that of the hydrophobic residues. Combining two features that decrease M_w,app_ do not lead to a further decrease but rather lead to a plateau at a ΔM_w,app_/residue of 90-100 Da.

**Figure 3:**
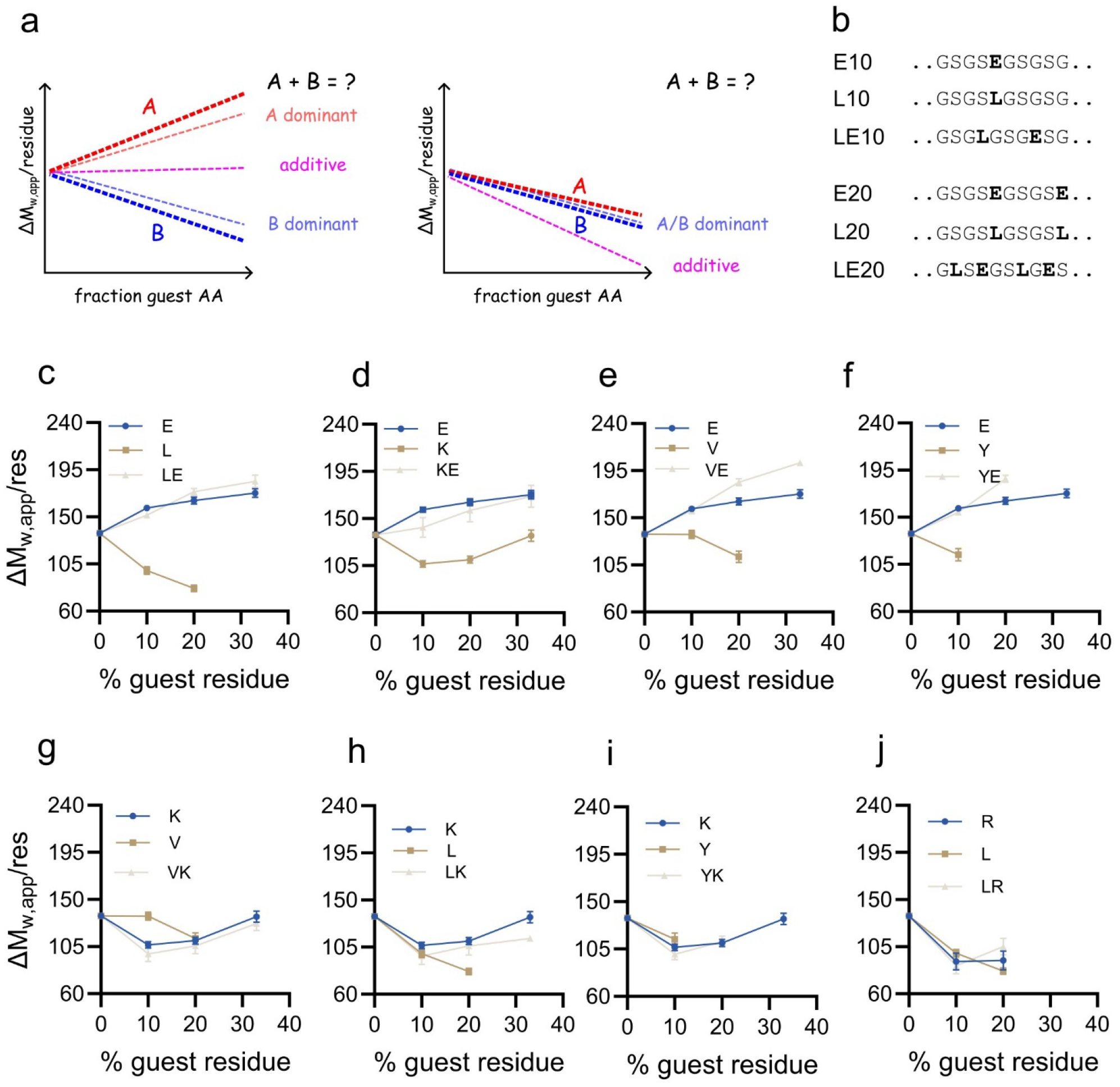
Additivity between individual sequence contributors to apparent molecular weight. a) Schematic representation of possible outcomes of combining guest residues that have opposing (left) or similar (right) effects on apparent molecular weight. Outcomes range from additivity to dominance of one factor. b) Sequence patterns used to assess the additivity of guest residues. c-j) ΔM_w,app_ of combinations of equal amounts of two guest residues compared to the ΔM_w,app_ of similar sequences with the individual residues. Error bars represent s.d. (n=3).

In total, our results confirm and extend our understanding of the sequence determinants of the high M_w,app_ of IDRs in SDS-PAGE. In agreement with previous studies, we found that the fraction of negatively charged residues is the strongest contributor to the elevated M_w,app_, likely due to poor binding of negatively charged sequences to negatively charged SDS micelles (Fig. 4). Similarly, we also show that neutral polar tracts have elevated M_w,app_. SDS can bind and denature proteins completely devoid of charges,^18^ however likely still require hydrophobic residues. Therefore, poor SDS binding can also explain the hypomobility of polar tracts such as the GS-repeats. Proline residues also increase M_w,app_ relative to a pure GS repeat, which is hard to ascribe to decreased SDS binding. Instead, proline residues lead to expansion of IDRs,^13,16^ which all-other-things-equal should lead to reduced electrophoretic mobility and an increase in M_w,app_. It thus suggests that both the compaction and SDS binding contribute to M_w,app_.

**Figure 4:**
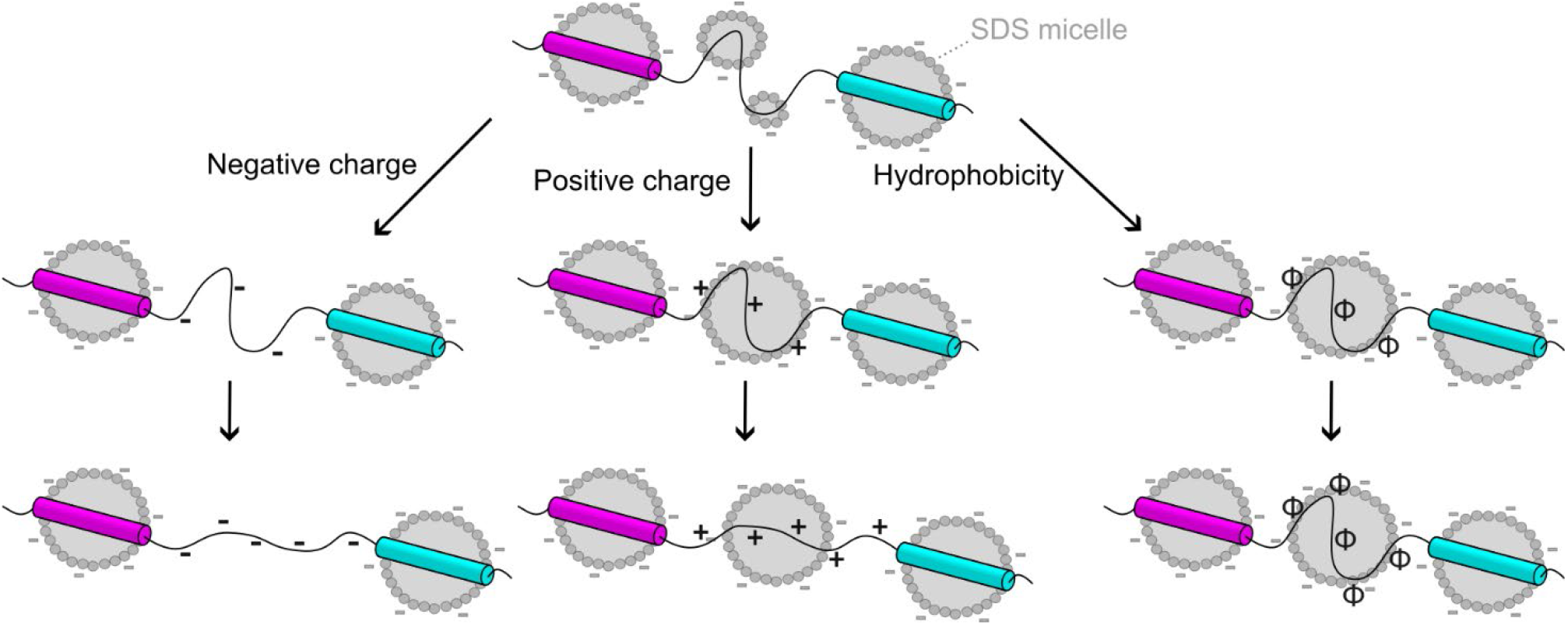
Schematic model for sequence effects on ΔM_w,app_ via the protein-decorated micelle model. Changes in IDR sequence can either lead to increased or decreased binding of SDS micelles, which translate into reduced or increased ΔM_w,app_, respectively. Beyond a certain point, further sequence changes do not lead to changes in the amount of bound SDS as segments are either fully bound or unbound. Beyond SDS binding, mobility is also affected by the sequence-dependent compaction of the IDRs.

Positively charged and hydrophobic residues initially decrease M_w,app_ in agreement with their expected effect on SDS binding (Fig. 4). However, lysine containing sequences showed a surprising biphasic effect where M_w,app_ increases at high charge densities. Comparison of K20 has a higher M_w,app_ than K10, whereas R20 is similar to R10. The expansion is thus not due to positive charge *per se* but is specific to lysine. Again, this can be understood by considering IDR compaction, where arginine residues generally induce compaction – e.g. as observed in identical synthetic IDR series as those tested here.^13^ The effect of positively charged guest residues thus also suggests that IDR compaction affects M_w,app_.

The simplest form type of sequence-based predictor of M_w,app_ of IDRs would simply tally up contributions from individual residues and combine them. Our test of additivity showed that this will not work: We neither observe additivity when combining features with opposite or similar effect on M_w,app_. The *protein-decorated micelle* model suggests an explanation for the lack of additivity as it suggests two extreme cases for the interaction of a protein with SDS: Fully bound or fully unbound. When an IDR segment is already fully bound to the micelle, increasing its affinity further would not increase SDS binding (Fig. 4). Similarly, it is plausible that an IDR rich in acidic residues cannot be induced to bind SDS by neither hydrophobicity nor compensating charged. This suggests that a hypothetical SDS-PAGE M_w,app_ predictor should consider the pattern and not just the total composition of residues in an IDR. Machine learning is well-suited to develop such a predictor, but requires large, structured data sets. Accordingly, the data acquired here in available in (Table S1, 2) to support such future efforts.

Especially the negatively charged IDRs were hard to detect in isolation – likely due to poor staining by the negatively charged Coomassie dye. Alternative staining strategies such as zinc or silver staining might be more appropriate for such IDRs in general. Here, we investigated the shift in SDS-PAGE mobility an IDR induces in a fusion protein (ΔM_w,app_) - in part as a proxy for the M_w,app_ of the isolated IDR. This strategy raises the question about whether ΔM_w,app_ is additive, e.g. whether addition of an IDR with M_w,app_ of 10 kDa result in ΔM_w,app_ of 10 kDa regardless of the fusion protein context. Comparison of the cleaved IDR in Fig. S1 (M_w,app_ = ~17 kDa) with the same IDR in a fusion protein (ΔM_w,app_ = ~22 kDa) suggests that sequence elements are not fully additive – although both values are noticeably higher than the true M_w_. The non-additivity can be explained by the dynamic structures of protein-detergent complexes formed in the presence of SDS (Fig. 4). Neighbouring regions may thus have positive or negative cooperativity in SDS binding, and proteins may form long-range contacts in the SDS-denatured state. Such effects would also need to be considered in developing an accurate predictor of SDS-PAGE mobility.

## Materials and methods

### Protein expression and purification

Codon optimized genes were purchased from Genscript cloned into the pET28a between the NdeI and XhoI sites. Proteins were expressed and purified using a generic high through-put protocol described previously.^19,20^ Before SDS-PAGE, purified proteins were dialyzed against 20 mM Na_2_HPO_4_/NaH_2_PO_4_ buffer (pH =7.5), 200 mM NaCl. Some fusion proteins were cleaved overnight at room temperature with 2 U/mg thrombin and subsequently loaded onto Superdex 75 Increase 10/300 GL column (Cytiva) equilibrated with 50 mM Na_2_HPO_4_/NaH_2_PO_4_ buffer (pH =7.5), 200 mM NaCl (Fig. S1).

### SDS-PAGE analysis

SDS-PAGE was conducted using mPAGE® 4-20% Bis-Tris precast gels (Millipore) run for 60 min at 120 V using a Pierce^TM^ unstained protein molecular weight marker (Ca. No. 26610, Thermo Scientific) as M_w_ reference. The amount of protein loaded into each well was optimized empirically to account for different staining efficiency. Gel loading order was varied between repeats, and the two outermost lanes were omitted to avoid distortions. Gels were digitalized using a Gel Doc^TM^ EZ Imager (Bio-Rad) and analysed using Image Lab V. 5.2.1 (Bio-Rad). Analysis was performed using the *automated lane detection* tool (with manual adjustments when needed) followed by the *band analysis function* and the automated calculation of M_w,app_ of by comparison to marker bands.

## Description of supplementary materials

The supplementary materials contain the sequences of proteins used, tabular values of M_w,app_ and uncropped scans of SDS-PAGE gels.

## Data availability

All data contained is included in the manuscript of supplementary materials.

## Acknowledgements

The work was supported by grants from the Novo Nordisk Foundation (NNF23OC0082071), the Danish National Research Foundation (DNRF133) and Independent Research Fund Denmark (2035-00285B).

