## Supplemental information for "Sequence determinants of the hypomobility of intrinsically disordered proteins in SDS-PAGE"

**Supplementary text:**

p. 2: Scheme of fusion protein design

**Supplementary tables:**

**Table S1:** Amino acid sequence of all the intrinsically disordered regions (IDRs) used in the study.

**Table S2:** Apparent molecular weights from SDS-PAGE of synthetic IDRs

**Supplementary figures:**

**Figure S1:** Size exclusion chromatography of IDRs released from coiled coils by proteolysis

**Figure S2:** Uncropped images of all the gels used in the main manuscript and for analysis

### Scheme of fusion protein design

**Sequence elements used in fusion proteins:**

Leader sequence from plasmid: MASMTGGQQMGRDP

His-tag: HHHHHH

Strep-tag: WSHPQFEK

MBD2 coiled coil: VTDEDIRKQEERVQQVRKKLEEALMAD

p66α coiled coil: PEERERMIKQLKEELRLEEAKLVLLKKLRQS

Thrombin cleavage site: LVPRGS

**General fusion protein sequence:**

MASMTGGQQMGRDPWSHPQFEKGSGSSVTDEDIRKQEERVQQVRKKLEEALMADILSGGGTLVPRGS

- (IDR insert)-

YLVPRGSEFGGPEERERMIKQLKEELRLEEAKLVLLKKLRQSQIQGGGGLEHHHHHH

**Example (120 residue GS repeat):**

MASMTGGQQMGRDPWSHPQFEKGSGSSVTDEDIRKQEERVQQVRKKLEEALMADILSGGGTLVPRGSGSGSGS
GSGSGSGSGSGSGSGSGSGSGSGSGSGSGSGSGSGSGSGSGSGSGSGSGSGSGSGSGSGSGSGSGSGSGSGSG
SGSGSGSGSGSGSGSGSGSGSGSGSGSGSGSGSGSGSGSGSGSGSYLVPRGSEFGGPEERERMIKQLKEELRLEEAK
LVLLKKLRQSQIQGGGGLEHHHHHH

**Table S1:** Amino acid sequence of all the intrinsically disordered regions (IDRs) used in the study

| Name | IDR sequence |
|---|---|
| **GS120** | GSGSGSGSGSGSGSGSGSGSGSGSGSGSGSGSGSGSGSGSGSGSGSGSGSGSGSGSGSGSGSGSGSG<br>SGSGSGSGSGSGSGSGSGSGSGSGSGSGSGSGSGSGSGSGSGSGSGSGSGSGSY |
| **E10** | GSGSEGSGSGSGSGESGSGSGSGSEGSGSGSGSGESGSGSGSGSEGSGSGSGSGESGSGSGSGSEGS<br>GSGSGSGESGSGSGSGSEGSGSGSGSGESGSGSGSGSEGSGSGSGSGESGSGSY |
| **E20** | GSGSEGSGSEGSGSEGSGSEGSGSEGSGSEGSGSEGSGSEGSGSEGSGSEGSGSEGSGSEGSGSEGS<br>GSEGSGSEGSGSEGSGSEGSGSEGSGSEGSGSEGSGSEGSGSEGSGSEGSGSEGSY |
| **E33** | GSEGSEGSEGSEGSEGSEGSEGSEGSEGSEGSEGSEGSEGSEGSEGSEGSEGSEGSEGSEGSEGSEG<br>SEGSEGSEGSEGSEGSEGSEGSEGSEGSEGSEGSEGSEGSEGSEGSEGSEGSEY |
| **K10** | GSGSKGSGSGSGSGKSGSGSGSGSKGSGSGSGSGKSGSGSGSGSKGSGSGSGSGKSGSGSGSGSKGS<br>GSGSGSGKSGSGSGSGSKGSGSGSGSGKSGSGSGSGSKGSGSGSGSGKSGSGSY |
| **K20** | GSGSKGSGSKGSGSKGSGSKGSGSKGSGSKGSGSKGSGSKGSGSKGSGSKGSGSKGSGSKGSGSKGS<br>GSKGSGSKGSGSKGSGSKGSGSKGSGSKGSGSKGSGSKGSGSKGSGSKGSGSKGSY |
| **K33** | GSKGSKGSKGSKGSKGSKGSKGSKGSKGSKGSKGSKGSKGSKGSKGSKGSKGSKGSKGSKGSKGSKG<br>SKGSKGSKGSKGSKGSKGSKGSKGSKGSKGSKGSKGSKGSKGSKGSKGSKGSKY |
| **R10** | GSGSRGSGSGSGSGRSGSGSGSGSRGSGSGSGSGRSGSGSGSGSRGSGSGSGSGRSGSGSGSGSRGS<br>GSGSGSGRSGSGSGSGSRGSGSGSGSGRSGSGSGSGSRGSGSGSGSGRSGSGSY |
| **R20** | GSRGSGSRGSGSRGSGSRGSGSRGSGSRGSGSRGSGSRGSGSRGSGSRGSGSRGSGSRGSGSRGSGS<br>RGSGSRGSGSRGSGSRGSGSRGSGSRGSGSRGSGSRGSGSRGSGSRGSGSRGSY |
| **D10** | GSGSDGSGSGSGSGDSGSGSGSGSDGSGSGSGSGDSGSGSGSGSDGSGSGSGSGDSGSGSGSGSDGS<br>GSGSGSGDSGSGSGSGSDGSGSGSGSGDSGSGSGSGSDGSGSGSGSGDSGSGSY |
| **L10** | GSGSLGSGSGSGSGLSGSGSGSGSLGSGSGSGSGLSGSGSGSGSLGSGSGSGSGLSGSGSGSGSLGS<br>GSGSGSGLSGSGSGSGSLGSGSGSGSGLSGSGSGSGSLGSGSGSGSGLSGSGSY |
| **L20** | GSLGSGSLGSGSLGSGSLGSGSLGSGSLGSGSLGSGSLGSGSLGSGSLGSGSLGSGSLGSGSLGSGS<br>LGSGSLGSGSLGSGSLGSGSLGSGSLGSGSLGSGSLGSGSLGSGSLGSGSLGSY |
| **V10** | GSGSVGSGSGSGSGVSGSGSGSGSVGSGSGSGSGVSGSGSGSGSVGSGSGSGSGVSGSGSGSGSVGS<br>GSGSGSGVSGSGSGSGSVGSGSGSGSGVSGSGSGSGSVGSGSGSGSGVSGSGSY |
| **V20** | GSVGSGSVGSGSVGSGSVGSGSVGSGSVGSGSVGSGSVGSGSVGSGSVGSGSVGSGSVGSGSVGSGS<br>VGSGSVGSGSVGSGSVGSGSVGSGSVGSGSVGSGSVGSGSVGSGSVGSGSVGSY |

| | |
|---|---|
| **P10** | GSPGSGSGSGSGPSGSGSGSGSPGSGSGSGSGPSGSGSGSGSPGSGSGSGSGPSGSGSGSGSPGSGSGSGSGSGPSGSGSGSGSPGSGSGSGSGPSGSGSGSGSPGSGSGSGSGPSGSGSY |
| **P20** | GSPGSGSPGSGSPGSGSPGSGSPGSGSPGSGSPGSGSPGSGSPGSGSPGSGSPGSGSPGSGSPGSGSPGSGSPGSGSPGSGSPGSGSPGSGSPGSGSPGSGSPGSGSPGSGSPGSGSPGSY |
| **P33** | GSPGSPGSPGSPGSPGSPGSPGSPGSPGSPGSPGSPGSPGSPGSPGSPGSPGSPGSPGSPGSPGSPGSPGSPGSPGSPGSPGSPGSPGSPGSPGSPGSPGSPGSPGSPGSPGSPGSPGSPY |
| **Y10** | GSGSYGSGSGSGSGYSGSGSGSGSYGSGSGSGSGYSGSGSGSGSYGSGSGSGSGYSGSGSGSGSYGSGSGSGSGYSGSGSGSGSYGSGSGSGSGYSGSGSGSGSYGSGSGSGSGYSGSGSY |
| **W10** | GSGSWGSGSGSGSGWSGSGSGSGSWGSGSGSGSGWSGSGSGSGSWGSGSGSGSGWSGSGSGSGSWGSGSGSGSGWSGSGSGSGSWGSGSGSGSGWSGSGSGSGSWGSGSGSGSGWSGSGSY |
| **VK10** | GSGVGSGKSGSGVGSGKSGSGSGVGSGKSGSGVGSGKSGSGSGVGSGKSGSGVGSGKSGSGSGVGSGKSGSGVGSGKSGSGSGVGSGKSGSGVGSGKSGSGSGVGSGKSGSGVGSGKSGSY |
| **VK20** | GVSKGSVGKSGVSKGSVGKSGVSKGSVGKSGVSKGSVGKSGVSKGSVGKSGVSKGSVGKSGVSKGSVGKSGVSKGSVGKSGVSKGSVGKSGVSKGSVGKSGVSKGSVGKSGVSKGSVGKSY |
| **VK33** | GVKSVKGVKSVKGVKSVKGVKSVKGVKSVKGVKSVKGVKSVKGVKSVKGVKSVKGVKSVKGVKSVKGVKSVKGVKSVKGVKSVKGVKSVKGVKSVKGVKSVKGVKSVKGVKSVKGVKSVKY |
| **LK10** | GSGLGSGKSGSGLGSGKSGSGSGLGSGKSGSGLGSGKSGSGSGLGSGKSGSGLGSGKSGSGSGLGSGKSGSGLGSGKSGSGSGLGSGKSGSGLGSGKSGSGSGLGSGKSGSGLGSGKSGSY |
| **LK20** | GLSKGSLGKSGLSKGSLGKSGLSKGSLGKSGLSKGSLGKSGLSKGSLGKSGLSKGSLGKSGLSKGSLGKSGLSKGSLGKSGLSKGSLGKSGLSKGSLGKSGLSKGSLGKSGLSKGSLGKSY |
| **LK33** | GLKSLKGLKSLKGLKSLKGLKSLKGLKSLKGLKSLKGLKSLKGLKSLKGLKSLKGLKSLKGLKSLKGLKSLKGLKSLKGLKSLKGLKSLKGLKSLKGLKSLKGLKSLKGLKSLKGLKSLKY |
| **VE10** | GSGVGSGESGSGVGSGESGSGSGVGSGESGSGVGSGESGSGSGVGSGESGSGVGSGESGSGSGVGSGESGSGVGSGESGSGSGVGSGESGSGVGSGESGSGSGVGSGESGSGVGSGESGSY |
| **VE20** | GVSEGSVGESGVSEGSVGESGVSEGSVGESGVSEGSVGESGVSEGSVGESGVSEGSVGESGVSEGSVGESGVSEGSVGESGVSEGSVGESGVSEGSVGESGVSEGSVGESGVSEGSVGESY |
| **VE33** | GVESVEGVESVEGVESVEGVESVEGVESVEGVESVEGVESVEGVESVEGVESVEGVESVEGVESVEGVESVEGVESVEGVESVEGVESVEGVESVEGVESVEGVESVEGVESVEGVESVEY |
| **LE10** | GSGLGSGESGSGLGSGESGSGSGLGSGESGSGLGSGESGSGSGLGSGESGSGLGSGESGSGSGLGSGESGSGLGSGESGSGSGLGSGESGSGLGSGESGSGSGLGSGESGSGLGSGESGSY |

| | |
|---|---|
| **LE20** | GLSEGSLGESGLSEGSLGESGLSEGSLGESGLSEGSLGESGLSEGSLGESGLSEGSLGESGLSEGSL<br>GESGLSEGSLGESGLSEGSLGESGLSEGSLGESGLSEGSLGESGLSEGSLGESY |
| **LE33** | GLESLEGLESLEGLESLEGLESLEGLESLEGLESLEGLESLEGLESLEGLESLEGLESLEGLESLEG<br>LESLEGLESLEGLESLEGLESLEGLESLEGLESLEGLESLEGLESLEGLESLEY |
| **YK10** | GSKGSGSYGSGSKGSGSYGSGSKGSGSYGSGSKGSGSYGSGSKGSGSYGSGSKGSGSYGSGSKGSGS<br>YGSGSKGSGSYGSGSKGSGSYGSGSKGSGSYGSGSKGSGSYGSGSKGSGSYGSY |
| **YK20** | GKSYGSKGYSGKSYGSKGYSGKSYGSKGYSGKSYGSKGYSGKSYGSKGYSGKSYGSKGYSGKSYGSK<br>GYSGKSYGSKGYSGKSYGSKGYSGKSYGSKGYSGKSYGSKGYSGKSYGSKGYSY |
| **YE10** | GSYGSGSEGSGSYGSGSEGSGSYGSGSEGSGSYGSGSEGSGSYGSGSEGSGSYGSGSEGSGSYGSGS<br>EGSGSYGSGSEGSGSYGSGSEGSGSYGSGSEGSGSYGSGSEGSGSYGSGSEGSY |
| **YE20** | GYSEGSYGESGYSEGSYGESGYSEGSYGESGYSEGSYGESGYSEGSYGESGYSEGSYGESGYSEGSY<br>GESGYSEGSYGESGYSEGSYGESGYSEGSYGESGYSEGSYGESGYSEGSYGESY |
| **GS120** | GSGSGSGSGSGSGSGSGSGSGSGSGSGSGSGSGSGSGSGSGSGSGSGSGSGSGSGSGSGSGSGSGSG<br>SGSGSGSGSGSGSGSGSGSGSGSGSGSGSGSGSGSGSGSGSGSGSGSGSGSGSY |
| **KE10** | GSKGSGSEGSGSKGSGSEGSGSKGSGSEGSGSKGSGSEGSGSKGSGSEGSGSKGSGSEGSGSKGSGS<br>EGSGSKGSGSEGSGSKGSGSEGSGSKGSGSEGSGSKGSGSEGSGSKGSGSEGSY |
| **KE20** | GKSEGSKGESGKSEGSKGESGKSEGSKGESGKSEGSKGESGKSEGSKGESGKSEGSKGESGKSEGSK<br>GESGKSEGSKGESGKSEGSKGESGKSEGSKGESGKSEGSKGESGKSEGSKGESY |
| **KE33** | GKESKEGKESKEGKESKEGKESKEGKESKEGKESKEGKESKEGKESKEGKESKEGKESKEGKESKEG<br>KESKEGKESKEGKESKEGKESKEGKESKEGKESKEGKESKEGKESKEGKESKEY |
| **KE50** | KEKEKEKEKEKEKEKEKEKEKEKEKEKEKEKEKEKEKEKEKEKEKEKEKEKEKEKEKEKEKEKEKEK<br>EKEKEKEKEKEKEKEKEKEKEKEKEKEKEKEKEKEKEKEKEKEKEKEKEKEKEY |
| **RL10** | GSGLGSGRSGSGLGSGRSGSGSGLGSGRSGSGLGSGRSGSGSGLGSGRSGSGLGSGRSGSGSGLGSG<br>RSGSGLGSGRSGSGSGLGSGRSGSGLGSGRSGSGSGLGSGRSGSGLGSGRSGSY |
| **RL20** | GLSRGSLGRSGLSRGSLGRSGLSRGSLGRSGLSRGSLGRSGLSRGSLGRSGLSRGSLGRSGLSRGSL<br>GRSGLSRGSLGRSGLSRGSLGRSGLSRGSLGRSGLSRGSLGRSGLSRGSLGRSY |

**Table S2:** Apparent molecular weights from SDS-PAGE of IDRs

| IDR | $M_{w,app}$ 1 | $M_{w,app}$ 2 | $M_{w,app}$ 3 | Av. $M_{w,app}$ | s.d. | $\Delta M_{w,app}$ | s.d. |
|---|---|---|---|---|---|---|---|
| **GS0** | 15.40 | 15.60 | 15.10 | 15.37 | 0.25 | 0.00 | 0.36 |
| **E10** | 33.70 | 33.60 | 34.00 | 33.77 | 0.21 | 18.40 | 0.33 |
| **E20** | 35.10 | 35.00 | 35.60 | 35.23 | 0.32 | 19.87 | 0.41 |
| **E33** | 35.90 | 35.80 | 36.60 | 36.10 | 0.44 | 20.73 | 0.50 |
| **K10** | 27.90 | 28.20 | 28.40 | 28.17 | 0.25 | 12.80 | 0.36 |
| **K20** | 28.40 | 28.60 | 29.00 | 28.67 | 0.31 | 13.30 | 0.40 |
| **K33** | 30.70 | 30.40 | 31.50 | 30.87 | 0.57 | 15.50 | 0.62 |
| **R10** | 26.40 | 25.30 | 27.10 | 26.27 | 0.91 | 10.90 | 0.94 |
| **R20** | 26.50 | 25.30 | 27.40 | 26.40 | 1.05 | 11.03 | 1.08 |
| **D10** | 33.70 | 33.80 | 34.70 | 34.07 | 0.55 | 18.70 | 0.61 |
| **L10** | 27.20 | 27.60 | 26.90 | 27.23 | 0.35 | 11.87 | 0.43 |
| **L20** | 25.10 | 25.20 | 24.70 | 25.00 | 0.26 | 9.63 | 0.37 |
| **V10** | 31.30 | 31.20 | 31.90 | 31.47 | 0.38 | 16.10 | 0.45 |
| **V20** | 28.50 | 28.20 | 29.30 | 28.67 | 0.57 | 13.30 | 0.62 |
| **Y10** | 28.90 | 28.50 | 29.80 | 29.07 | 0.67 | 13.70 | 0.71 |
| **W10** | 25.60 | 25.20 | 26.80 | 25.87 | 0.83 | 10.50 | 0.87 |
| **P10** | 32.10 | 32.00 | 31.80 | 31.97 | 0.15 | 16.60 | 0.29 |
| **P20** | 32.80 | 32.80 | 32.20 | 32.60 | 0.35 | 17.23 | 0.43 |
| **P33** | 33.40 | 33.60 | 32.90 | 33.30 | 0.36 | 17.93 | 0.44 |
| **RL20** | 27.90 | 26.80 | 28.60 | 27.77 | 0.91 | 12.40 | 0.94 |
| **RL10** | 25.80 | 24.90 | 26.70 | 25.80 | 0.90 | 10.43 | 0.93 |
| **VK33** | 29.80 | 31.30 | 30.70 | 30.60 | 0.75 | 15.23 | 0.80 |
| **VK20** | 28.20 | 27.10 | 28.80 | 28.03 | 0.86 | 12.67 | 0.90 |
| **VK10** | 27.40 | 26.20 | 27.80 | 27.13 | 0.83 | 11.77 | 0.87 |
| **LK33** | 28.80 | 29.10 | 29.10 | 29.00 | 0.17 | 13.63 | 0.31 |
| **LK20** | 28.60 | 27.20 | 28.50 | 28.10 | 0.78 | 12.73 | 0.82 |
| **LK10** | 27.60 | 26.30 | 27.40 | 27.10 | 0.70 | 11.73 | 0.74 |
| **VE33** | 39.90 | 39.50 | 39.60 | 39.67 | 0.21 | 24.30 | 0.33 |
| **VE20** | 37.70 | 37.50 | 37.10 | 37.43 | 0.31 | 22.07 | 0.40 |
| **VE10** | 34.30 | 34.50 | 33.90 | 34.23 | 0.31 | 18.87 | 0.40 |
| **LE33** | 37.40 | 38.10 | 36.80 | 37.43 | 0.65 | 22.07 | 0.70 |
| **LE20** | 35.80 | 36.20 | 35.70 | 35.90 | 0.26 | 20.53 | 0.37 |
| **LE10** | 33.30 | 33.20 | 33.30 | 33.27 | 0.06 | 17.90 | 0.26 |
| **KE50** | 41.80 | 41.40 | 43.20 | 42.13 | 0.95 | 26.77 | 0.98 |
| **KE33** | 35.40 | 35.00 | 37.30 | 35.90 | 1.23 | 20.53 | 1.25 |
| **KE20** | 33.60 | 33.50 | 35.80 | 34.30 | 1.30 | 18.93 | 1.32 |
| **KE10** | 31.50 | 31.90 | 33.60 | 32.33 | 1.12 | 16.97 | 1.14 |
| **GS120** | 31.50 | 31.30 | 31.70 | 31.50 | 0.20 | 16.13 | 0.32 |

| | | | | | | | |
|---|---|---|---|---|---|---|---|
| **YE10** | 33.80 | 34.00 | 34.00 | 33.93 | 0.12 | 18.57 | 0.28 |
| **YE20** | 37.90 | 38.00 | 37.40 | 37.77 | 0.32 | 22.40 | 0.41 |
| **YK10** | 26.80 | 27.40 | 27.90 | 27.37 | 0.55 | 12.00 | 0.61 |
| **YK20** | 28.20 | 29.30 | 28.90 | 28.80 | 0.56 | 13.43 | 0.61 |
| **WE10** | 30.90 | 33.10 | 32.10 | 32.03 | 1.10 | 16.67 | 1.13 |
| **WK10** | 26.20 | 28.40 | 27.20 | 27.27 | 1.10 | 11.90 | 1.13 |

#### Supporting Figures

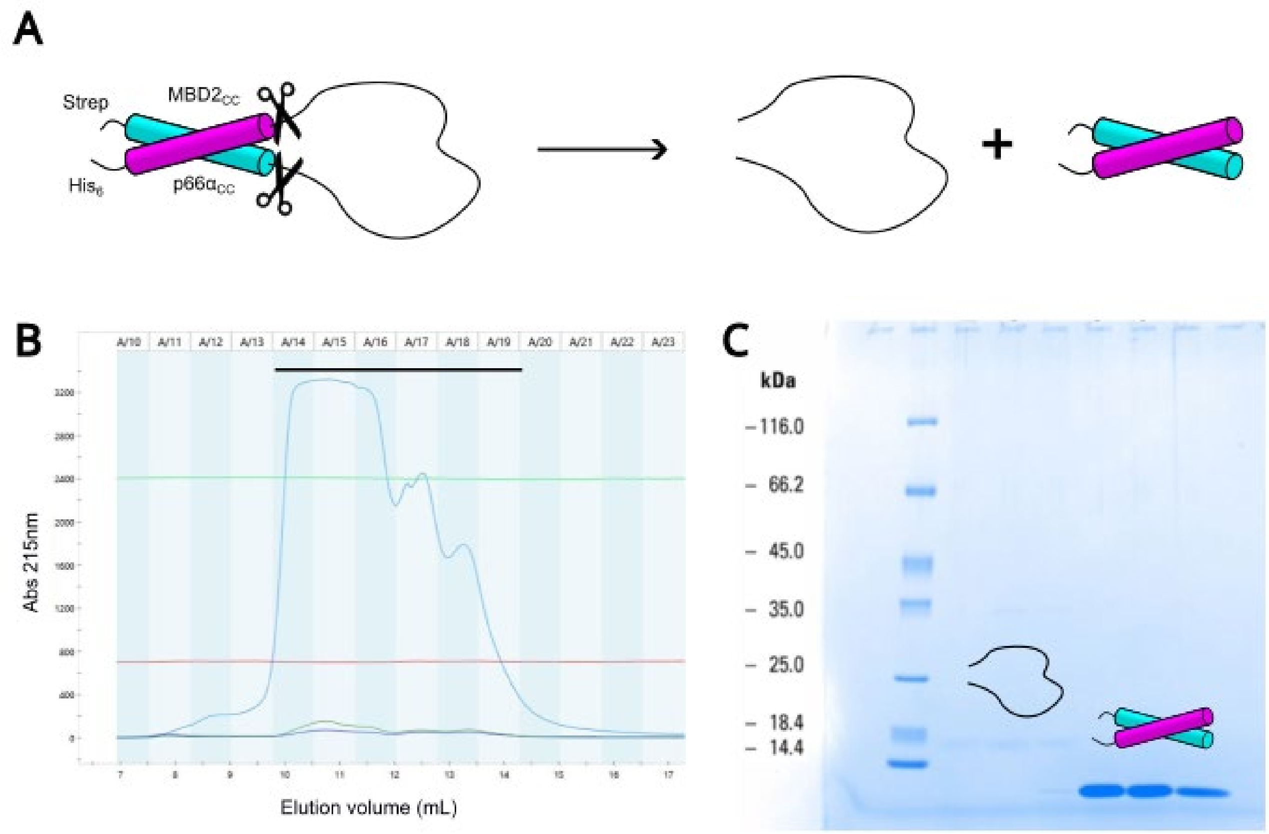


**Figure S1: Size exclusion chromatography of IDRs released from coiled coils by proteolysis.** (A) the isolated IDR can be released from the fusion protein by thrombin cleavage resulting in mixture of the free coiled-coils and the IDR. (B) Purification of the VE20 IDR after cleavage by size exclusion chromatography shows a major peak followed by smaller peaks. The bar indicates the six fractions selected for SDS-PAGE analysis. (C) SDS-PAGE of SEC fractions show that the major peak corresponds to free IDR, which is only faintly visible at an $M_{w,app}$ ~16 kDa, whereas the coiled-coil helices elute in the second peak and are clearly visible on the gel at $M_{w,app} < 10$ kDa.

**Figure S2:** Uncropped images of all the gels used in the main manuscript and for analysis

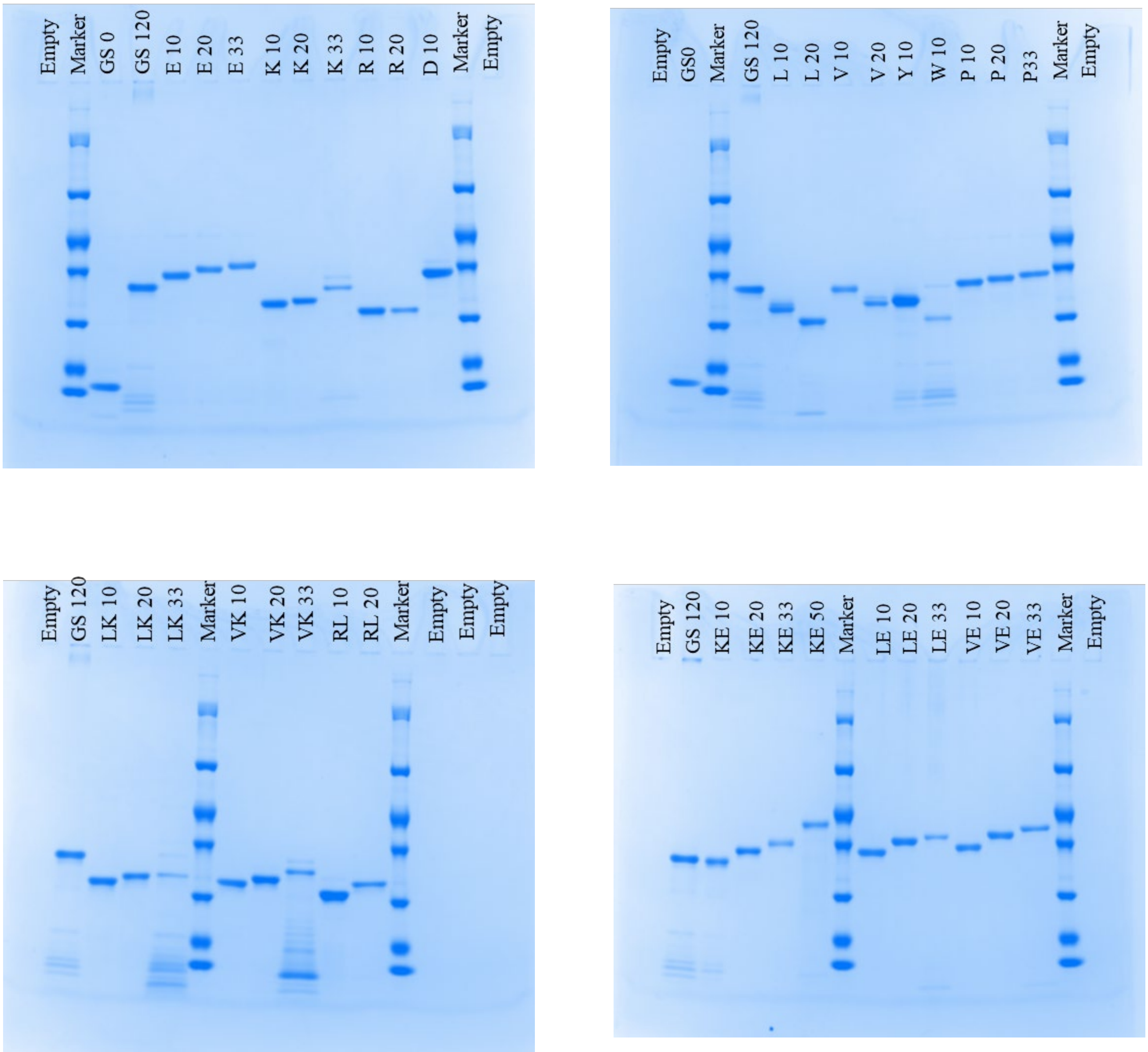

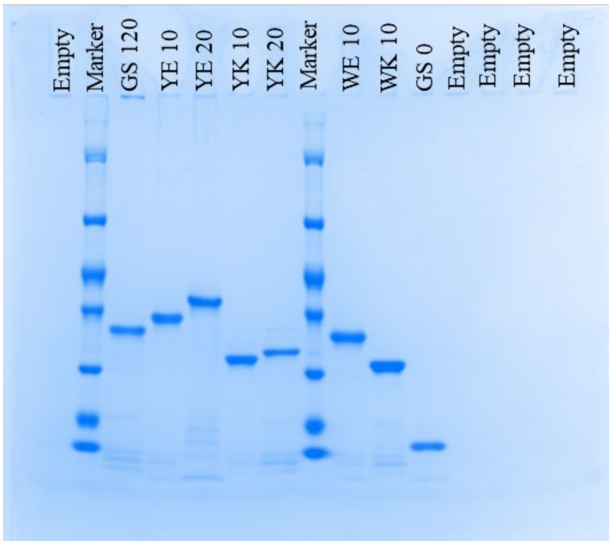
Empty
Marker
GS 120
YE 10
YE 20
YK 10
YK 20
Marker
WE 10
WK 10
GS 0
Empty
Empty
Empty
Empty

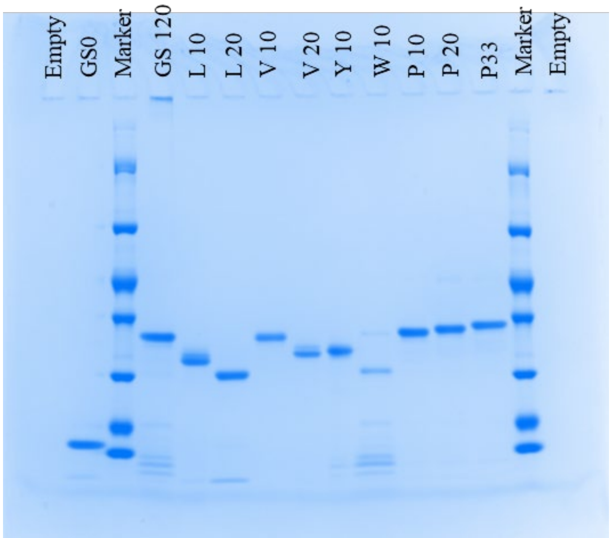
Empty
GS0
Marker
GS 120
L 10
L 20
V 10
V 20
Y 10
W 10
P 10
P 20
P33
Marker
Empty

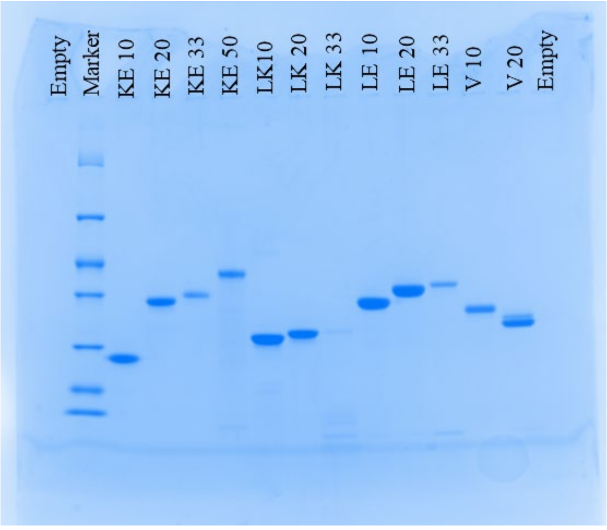
Empty
Marker
KE 10
KE 20
KE 33
KE 50
LK10
LK 20
LK 33
LE 10
LE 20
LE 33
V 10
V 20
Empty

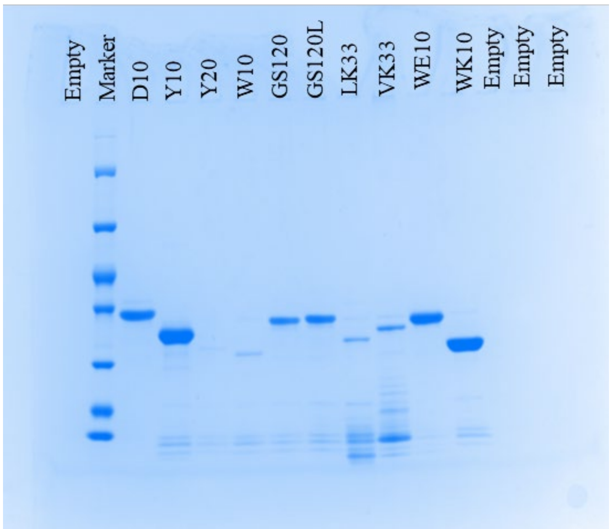
Empty
Marker
D10
Y10
Y20
W10
GS120
GS120L
LK33
VK33
WE10
WK10
Empty
Empty
Empty

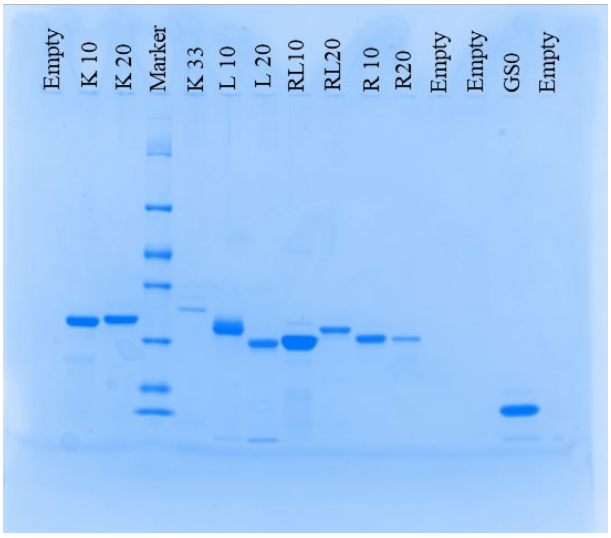
Empty
K 10
K 20
Marker
K 33
L 10
L 20
RL10
RL20
R 10
R20
Empty
Empty
GS0
Empty

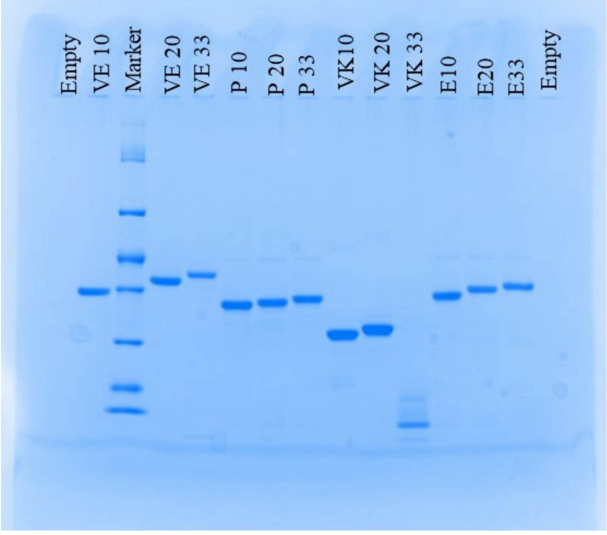
Empty
VE 10
Marker
VE 20
VE 33
P 10
P 20
P 33
VK10
VK 20
VK 33
E10
E20
E33
Empty

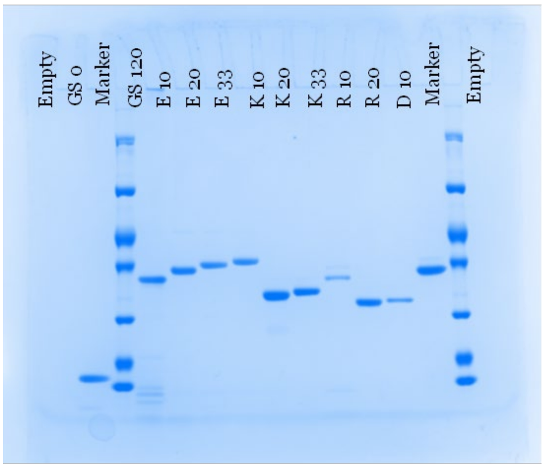
Empty
GS 0
Marker
GS 120
E 10
E 20
E 33
K 10
K 20
K 33
R 10
R 20
D 10
Marker
Empty
